# Opportunistic interactions on Fe^0^ between methanogens and acetogens from a climate lake

**DOI:** 10.1101/556704

**Authors:** Paola Andrea Palacios, Amelia-Elena Rotaru

**Affiliations:** Nordcee, Department of Biology, University of Southern Denmark, Odense, Denmark

**Keywords:** microbial influenced corrosion, acetogens, methanogens, interspecies interactions, iron corrosion, Clostridium, Methanosarcinales, Methanothermobacter

## Abstract

Microbial-induced corrosion has been extensively studied in pure cultures. However, Fe^0^ corrosion by complex environmental communities, and especially the interplay between microbial physiological groups, is still poorly understood. In this study, we combined experimental physiology and metagenomics to explore Fe^0^-dependent microbial interactions between physiological groups enriched from anoxic climate lake sediments. Then, we investigated how each physiological group interacts with Fe^0^. We offer evidence for a new interspecies interaction during Fe^0^ corrosion. We showed that acetogens enhanced methanogenesis but were negatively impacted by methanogens (opportunistic microbial interaction). Methanogens were positively impacted by acetogens. In the metagenome of the corrosive community, the acetogens were mostly represented by *Clostridium* and *Eubacterium,* the methanogens by *Methanosarcinales*, *Methanothermobacter* and *Methanobrevibacter.* Within the corrosive community, acetogens and methanogens produced acetate and methane concurrently, however at rates that cannot be explained by abiotic H_2_-buildup at the Fe^0^ surface. Thus, microbial-induced corrosion might have occurred via a direct or enzymatically mediated electron uptake from Fe^0^. The shotgun metagenome of *Clostridium* within the corrosive community contained several H_2_-releasing enzymes including [FeFe]-hydrogenases, which could boost Fe^0^-dependent H_2_-formation as previously shown for pure culture acetogens. Outside the cell, acetogenic hydrogenases could become a common good for any H_2_/CO_2_-consuming member in the microbial community including methanogens that rely on Fe^0^ as a sole energy source. However, the exact electron uptake mechanism from Fe^0^ remains to be unraveled.

## 1 Introduction

Steel infrastructure extends for billions of kilometers below ground enabling transport and storage of clean water, chemicals, fuels, sewage, but also protection for telecommunication and electricity cables. Climate change has led to extreme weather conditions like severe storms and rainfall. Urban storm and rainfall management in many countries especially northers countries like Denmark involves so called climate lakes (also known as storm water ponds or retention ponds) harvesting rainfall at large scale thus alleviating stormwater runoff in the cities (Mishra et al., 2020). If stormwater runoff is improperly detained, undergrown steel infrastructure could suffer tremendous damage. Damages induced by microbial induced corrosion (MIC) in the underground are often discovered too late, leading to environmental and economic devastation. Thus, it is important to be able to predict the lifespan of the material if exposed to microbial communities native to the site where steel structures are located. This would lead to effective replacement strategies and recuperation of the metal prior to accidental spills that may be detrimental to the surrounding environment (Arriba-Rodriguez et al., 2018; Skovhus et al., 2017; Usher et al., 2014a).

Corrosion occurring in climate lakes is poorly understood. In this study, we investigate corrosion by microorganisms from the anoxic sediments of a danish climate lake. In such anoxic environments where non-sulfidic conditions prevail, steel was expected to persist unharmed for centuries (Arriba-Rodriguez et al., 2018; Skovhus et al., 2017; Usher et al., 2014a). And yet, researchers showed that certain groups of anaerobes (methanogens and acetogens) strip electrons off the Fe^0^ surface leading to MIC (Kato et al., 2015; Mand et al., 2014, 2016; Mori et al., 2010; Zhang et al., 2003). Previous studies showed that MIC in non-sulfidic environments is often linked to the presence of acetogens like *Clostridium* and methanogens like *Methanosarcinales* on the surface of the corroded steel structure (Kato et al., 2015; Mand et al., 2014, 2016; Mori et al., 2010; Zhang et al., 2003; Zhu et al., 2003). It was therefore suggested that *Methanosarcinales* were indirectly involved in corrosion, growing in a mutualistic relationship with the acetogens, and allegedly both groups were gaining from the interaction (Mand et al., 2016; Zhang et al., 2003). This assumption was based on acetogens producing acetate, which would be then consumed by acetotrophic *Methanosarcinales* methanogens, acetogens were expected to be favored by the removal of their metabolic product - acetate. Such a mutualistic association on Fe^0^ between acetogenic *Clostridium* and methanogenic *Methanosarcinales* has not been yet demonstrated. In contrast, we recently showed that instead of acting cooperatively on Fe^0^, acetogens and methanogens competed for Fe^0^-electrons (Palacios et al., 2019a). Since, interspecies interactions on Fe^0^ are scantily examined, here we investigated the interplay between climate lake acetogens (reaction 1) and methanogens (reaction 2) when provided solely with Fe^0^ as their electron donor.

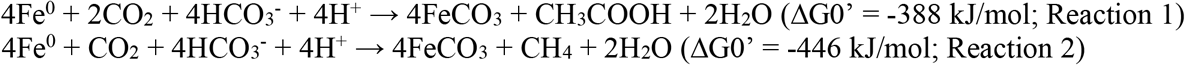

Theoretically, under standard thermodynamic conditions, methanogens should have an advantage over acetogens when provided with Fe^0^ as sole electron donor (Reactions 1 & 2). Especially, since methanogens, unlike acetogens, are more effective at retrieving abiotic H_2_ (formed on Fe^0^) due to their low H_2_-uptake thresholds (Drake et al., 2002; Kotsyurbenko et al., 2001). Several groups of methanogens could corrode Fe^0^ independent of acetogenic bacteria, including species of *Methanosarcina* (Belay and Daniels, 1990; Boopathy and Daniels, 1991; Daniels et al., 1987), *Methanobacterium* (Belay and Daniels, 1990; Dinh et al., 2004; Lorowitz et al., 1992) and *Methanococcus* (Lienemann et al., 2018c; Mori et al., 2010; Tsurumaru et al., 2018; Uchiyama et al., 2010). The mechanism by which methanogens corrode Fe^0^, has been debated and includes reports which suggest they retrieve abiotic-H_2_ off the Fe^0^ surface (Daniels et al., 1987), retrieve electrons directly using an unknown electron-uptake mechanism (Beese-Vasbender et al., 2015; Dinh et al., 2004) or use extracellular enzymes, which stimulate enzymatic H_2_-evolution or formate generation on the Fe^0^-surface (Deutzmann et al., 2015a; Lienemann et al., 2018c; Tsurumaru et al., 2018). The later mechanisms were especially relevant for *Methanococcus* species which harbored an unstable genomic island encoding [NiFe]-hydrogenases along with the heterodisulfide reductase super complex involved in formate generation (Lienemann et al., 2018c; Tsurumaru et al., 2018). And yet, oftentimes acetogens dominate corrosive communities, outcompeting methanogens when concentrations of H_2_ are high and temperatures are low, presumably due to the higher kinetics (V_max_) of their hydrogenases (Kotsyurbenko et al., 2001). Moreover, unlike methanogens, acetogens contain [FeFe]-hydrogenases (Peters et al., 2015), which could retrieve electrons directly from Fe^0^ for proton reduction to H_2_ (Mehanna et al., 2008, 2016; Rouvre and Basseguy, 2016).

In this study, we were interested to understand the interspecies dynamics on Fe^0^ of acetogens and methanogens from a climate lake. We used a combination of physiological experiments, inhibition strategies and whole metagenomic analyses to study the interactions of acetogens and methanogens during Fe^0^ corrosion. In contrast to our previous report on a corrosive coastal community (Palacios et al., 2019a), in this climate lake we observed a different type of microbial interaction, where acetogens (*Clostridium*) were negatively impacted by the presence of methanogens whereas methanogens were benefiting from their presence in a parasitic (−/+) type of interaction while inducing Fe^0^ corrosion.

## 2 Materials and methods

### 2.1 Sample collection and enrichment culture conditions

Sediment cores were sampled during the month of July 2016 from a small climate lake located near a construction site on the campus of the University of Southern Denmark (SDU), Odense. The salinity of the lake was 0.6 psu, and gas bubbles (including methane) were continuously released to the water surface while sampling. Sediment cores were sliced in the laboratory, and the depth horizon 15-20 cm was used for downstream enrichments in a DSM modified 120 media (modifications: 0.6g/L NaCl, without casitone, without sodium acetate, without methanol, and without Na_2_S × 9H_2_O). The enrichment cultures were prepared in 50 mL blue butyl-rubber-stoppered glass vials with an anoxic headspace of a CO_2_: N_2_ gas mix (20:80, v/v). Iron granules (99.98% Thermo Fisher, Germany) or iron coupons (3cm × 1cm × 1mm) were the only source of electrons over the course of five successive transfers. All incubations were performed in triplicate.

All enrichments were transferred when methane production reached stationary phase. DNA extractions, SEM analyses, and further experiments were performed at the fourth transfer, after 2 years of enrichment on Fe^0^. In addition, methanogen-specific coenzyme F_420_ auto-fluorescence was monitored via routine microscopy to confirm the presence or absence of methanogens. To evaluate the solitary corrosive potential of methanogens, we blocked all bacteria with an antibiotic cocktail 200 μg/mL of kanamycin and 100 μg/mL of ampicillin as done before (Palacios et al., 2019a). To evaluate the solitary corrosive potential of the acetogens, we inhibited all methanogens with 2 mM 2-bromoethanesulfonate (BES) (Zhou et al., 2011).

### 2.2 Chemical analyses

Methane and hydrogen concentrations were analyzed on a Thermo Scientific Trace 1300 gas chromatograph system coupled to a thermal conductivity detector (TCD). The injector was operated at 150°C and the detector at 200°C with 1.0 mL/min argon as reference gas. The oven temperature was constant at 70°C. Separation was done on a TG-BOND Msieve 5A column (Thermo Scientific; 30-m length, 0.53-mm i.d., and 20-μm film thickness) with argon as carrier gas at a flow of 25 mL/min. The GC was controlled and automated by a Chromeleon software (Dionex, Version 7). On our set-up the limit of detection for H_2_ and CH_4_ was 5 μM.

Acetate production was measured using the Dionex ICS-1500 Ion Chromatography System (ICS-1500) equipped with the AS50 autosampler, and an IonPac AS22 column coupled to a conductivity detector (31 mA). For separation of volatile fatty acids, we used 4.5 mM Na_2_CO_3_ with 1.4 mM NaHCO_3_ as eluent. The run was isothermic at 30°C with a flow rate of 1.2mL/min. The limit of detection for acetate was 0.1 mM.

### 2.3 DNA purification and metagenomic analyses

DNA was isolated as previously described (Palacios et al., 2019a), using a combination of two commercially available kits: MasterPure™ Complete DNA and RNA Purification Kit (Epicenter, Madison, Wi, USA), and the Fast Prep spin MP_tm_ kit for soil (Mobio/Qiagen, Hildesheim, Germany). DNA quality was verified on an agarose gel, and DNA was quantified on a mySPEC spectrophotometer (VWR**®**/ Germany). Whole metagenome sequencing was performed on a NovaSeq 6000 system, using an Illumina TrueSeq PCR-free approach via a commercially available service (Macrogen/ Europe). Unassembled DNA sequences were merged, quality checked, and annotated using the Metagenomics Rapid Annotation (MG-RAST) server (v4.03) with default parameters (Meyer et al., 2008). Illumina True Seq sequencing resulted in 1,982,966 high-quality reads of a total of 2,137,679 with an average length of 102 bp. For taxonomic analyses, the metagenomic data was compared with the RefSeq (Tatusova et al., 2015) database available on the MG-RAST platform. The rarefaction curve indicated that most of the prokaryotic diversity was covered in our sample. To investigate functional genes in the metagenome, sequencing reads were annotated against the KEGG Orthology (KO) reference database. Both taxonomic and functional analyses were performed with the following cutoff parameters: *e*-value of 1e–5, a minimum identity of 80%, and a maximum alignment length of 15 bp. The metagenome data is available at MG-RAST with this ID:xxxxxx.

### 2.4 Removal of corrosion crust and corrosion rates

The corrosion crust from the iron coupons was removed with inactivated acid (10% hexamine in 2M HCl) (Enning and Garrelfs, 2014). Then, the iron coupons were dried with N_2_ gas stream, and weighted.

### 2.5 Scanning electron microscopy

Fixation of cells on iron coupons was performed anaerobically by adding 2.5% glutaraldehyde in 0.1M phosphate buffer (pH 7.3) and incubating at 4°C for 12 h. The corroded coupons were then washed three times with 0.1 M phosphate buffer at 4°C for 10 min each. Dehydration was accomplished by a series of anoxic pure ethanol steps (each step 10 min; 35%, 50%, 70%, 80%, 90%, 95% and 100% v/v) (Araujo et al., 2003). The coupons were chemical dried with hexamethyldisilazane under a gentle N_2_ gas stream. Specimens were stored under N_2_ atmosphere and analyzed within 18-24 h at the UMASS electron microcopy facility using the FEI Magellan 400 XHR-SEM with a resolution of 5kV.

## 3 Results and discussion

Here we report on an opportunistic (+/−) interaction on Fe^0^, between methanogens and acetogens from the sediments of a climate lake. In the absence of any other terminal electron acceptor but CO_2_, microorganisms are expected to corrode Fe^0^ via either CO_2_-reductive methanogenesis (CO_2_ + 8e^−^ + 8H^+^ → CH_4_ + 2H_2_O / CO_2_-reductive methanogenesis) or via acetogenesis coupled with acetoclastic methanogenesis (2CO_2_ + 8e^−^ + 8H^+^ → CH_3_COOH + 2H_2_O / acetogenesis; CH_3_COOH → CH_4_ + CO_2_ / acetoclastic methanogenesis).

For the later, acetoclastic methanogens were commonly assumed to be indirectly involved in the corrosion of Fe^0^ since they were expected to establish a mutualistic association (+/+) with acetogens (Mand et al., 2016; Zhang et al., 2003). However, this is not always the case. We recently showed that coastal marine methanogens competed (−/−) with acetogens for Fe^0^-electrons (Palacios et al., 2019b). But now, we bring evidence for an opportunistic association between methanogens and acetogens from the sediments of a climate-lake. The methanogens from these sediments were favored by the acetogens however could not be explained by acetate-consumption alone. Acetogenic [FeFe] hydrogenases were heavily represented in the acetogens’ metagenome and may induce significant electron uptake from Fe^0^ coupled with H_2_-production. When released, hydrogenases may indiscriminately stimulate the growth of opportunistic hydrogenotrophic methanogens.

### 3.1 Pitted Fe^0^ corrosion

We monitored corrosion by a climate lake community maintained for 2 years on Fe^0^ in the absence of any typical electron acceptors. A time course of corrosion was taken during the fourth transfer on Fe^0^. The microbial community corroded Fe^0^ significantly in contrast to cell-free controls according to microscopy observations (Fig. 1), gravimetric measurements (Fig. 2) and metabolic product build-up (Fig. 3).

**Figure 1.**
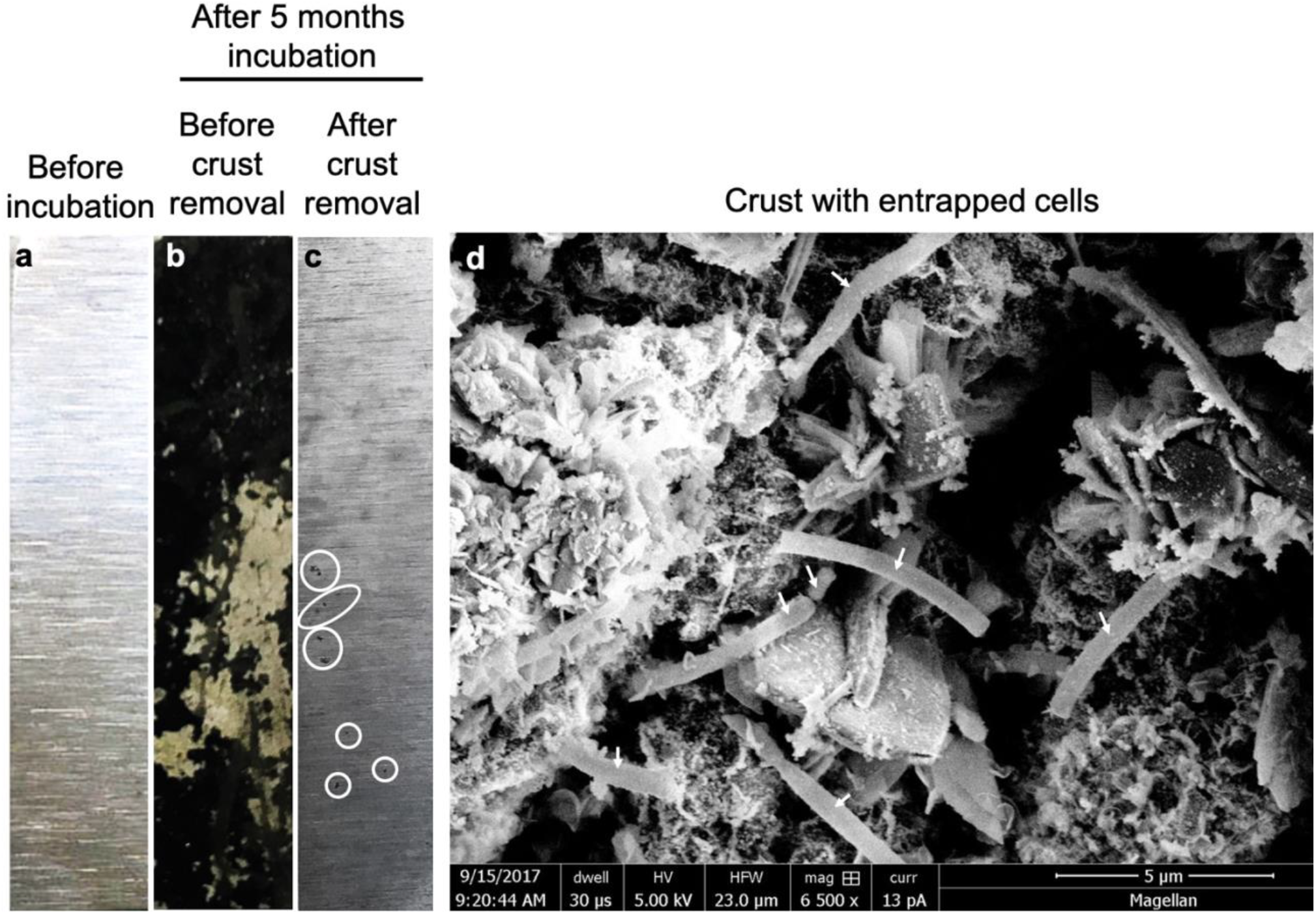
Pitted corrosion by microorganisms. Metallic iron sheets prior to exposure to cells (a), prior to removal of the crust (b) and after crust removal. (d) A scanning electron micrograph showing cells entrenched on the metal surface.

**Figure 2.**
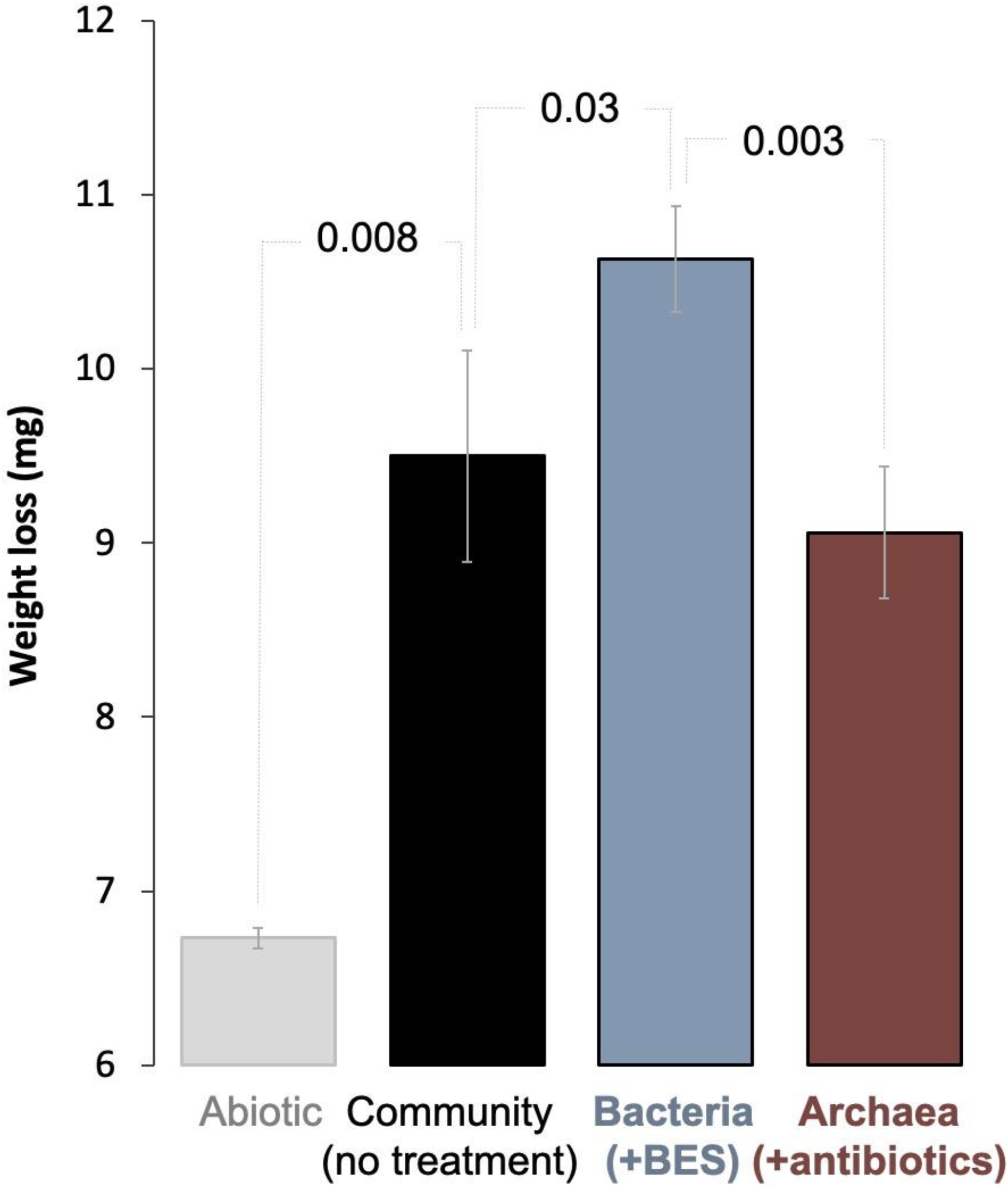
**Gravimetric determination of Fe^0^ corrosion** in abiotic controls (gray), by a microbial community from SDU sediments (black), by bacteria alone, after specific inhibition of the methanogens with 2-bromoethanesulfonate (blue), and methanogens alone after inhibition of all bacteria using a mix of antibiotics (red). n=3, p<0.05.

**Figure 3.**
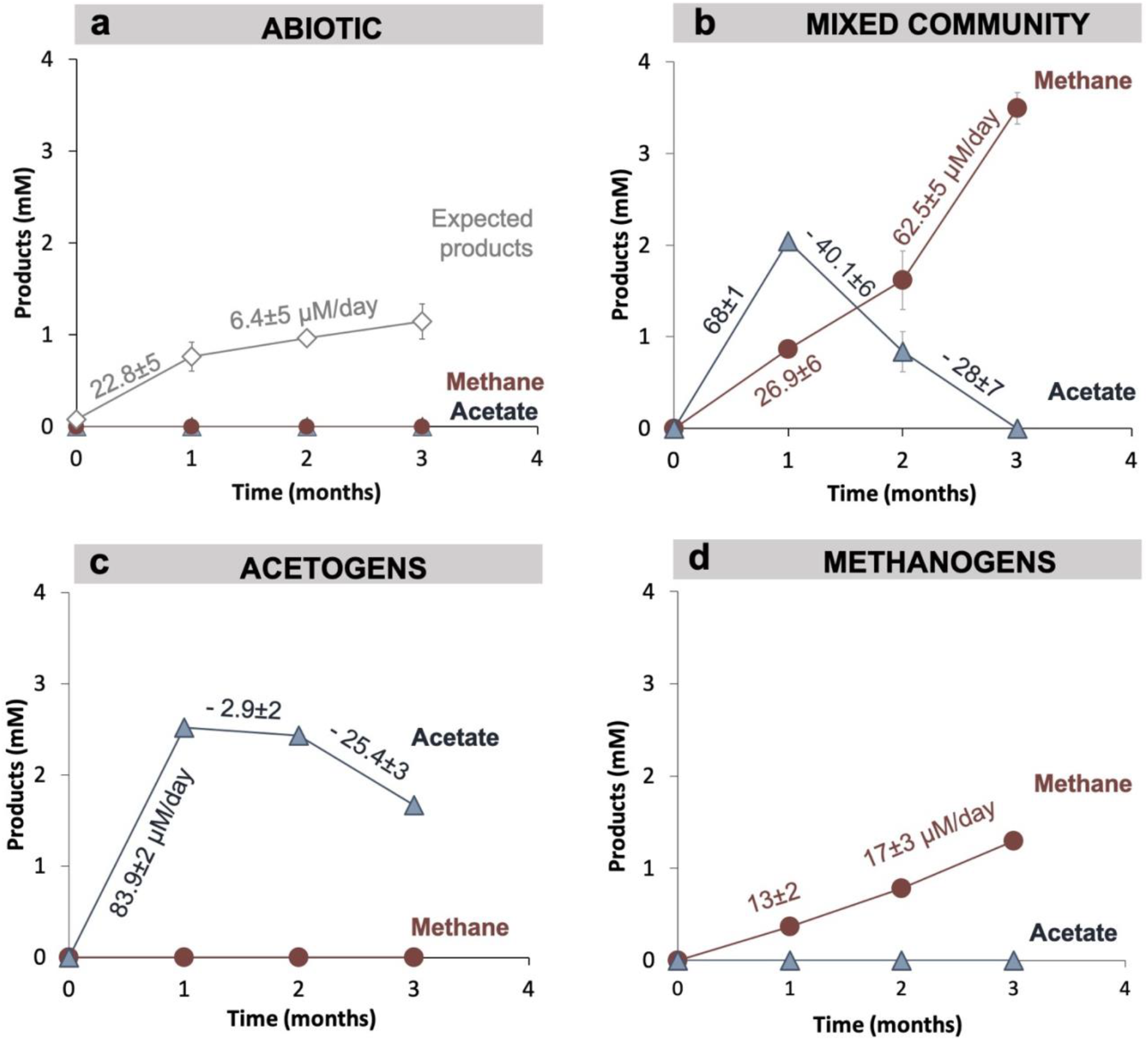
**Product formation** (methane – red; acetate – blue) by (a) abiotic controls (none), (b) a mixed acetogenic-methanogenic community; (c) unaccompanied acetogens, obtained by the specific inhibition of methanogens with 2-bromoethanesulfonate; (d) unaccompanied methanogens, obtained by specific inhibition of all bacteria with an antibiotic mix of kanamycin and ampicillin. All incubations were run in parallel and in triplicate (n=3).

After incubation for several months in the presence of cells the iron sheets were coated with a black crust, which after removal revealed small pitted corrosion underneath (Fig. 1). Scanning electron microscopy prior to crust removal shows cells entrapped on Fe^0^ in between minerals likely formed due to biocorrosion (Fig. 1d).

Gravimetric measurements showed that when this climate lake community was provided with Fe^0^ it induced 41% more weight loss and consumed 2.8 ± 0.6 mg more Fe^0^ (n=3; p=0.008) than cell-free controls (Fig. 2).

Metabolic product build-up showed that the microbial-community generated methane and acetate simultaneously once provided with Fe^0^ (Fig. 3), confirming that Fe^0^ delivers electrons for two types of microbial metabolisms, acetogenesis and methanogenesis. As expected, the abiotic controls with Fe^0^ showed no traces of microbial metabolic products (methane and acetate) (Fig. 3a).

During 3-months long incubations with Fe^0^, the corrosive community generated acetate transiently (first month), and ultimately accumulated only methane (Fig. 3b). At the start of the incubation (first month), acetogenesis rates (68±1 μM acetate/day) surpassed methanogenesis rates (27±6 μM methane/day) (Fig. 3, Fig. 4a). Whereas during the last two months of incubation, acetogenesis ceased. At the same time, methanogenesis sped up achieving rates twofold (62.5±5.1 μM methane/day) above those predicted (28±7.3 μM methane/day) by acetoclastic methanogenesis (Fig. 3, Fig. 4a). These results show that methanogens did not rely on the acetate generated by acetogens for methanogenesis. Altogether, the microbial community produced 3.3-fold more methane (3.5 ± 0.1 mM) than expected (1.1±0.2 mM) from the H_2_ evolved abiotically (2e^−^ + 2H^+^ → H_2_) at the Fe^0^ surface (Fig. 5). These results show that the microbial community employs effective alternative mechanisms (other than abiotic H_2_) to access electrons from the Fe^0^-surface. Unraveling these mechanisms is difficult without pure cultures. Nevertheless, we attempted to determine how the microorganisms influence each other’s metabolism and Fe^0^-corrosion by separating each physiologic group with group-specific inhibitors.

**Figure 4.**
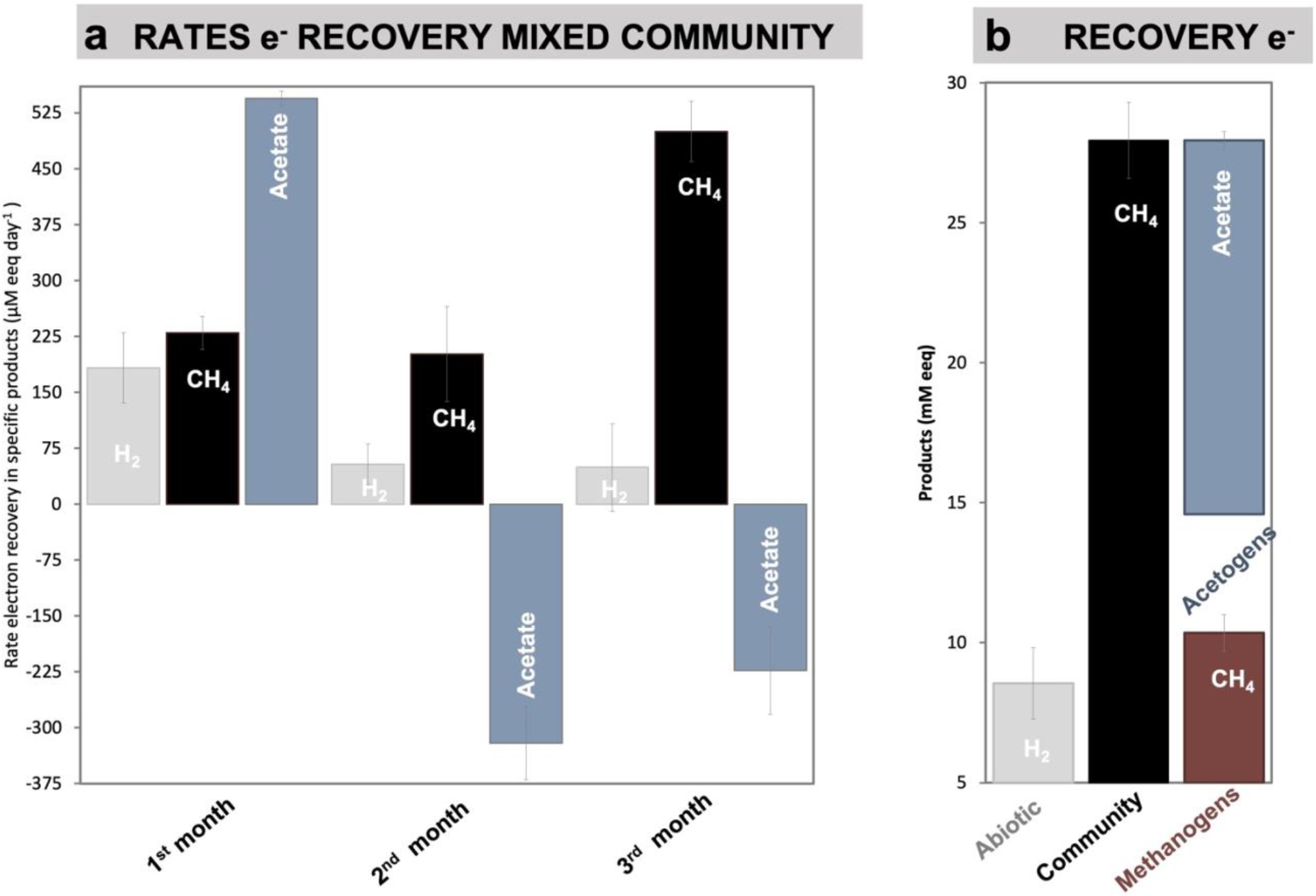
Electron recovery. Changes in electron recovery rates over the course of three months (a) and total electron recovery into products as mM electron equivalents (eeq) produced from Fe^0^ under four different conditions (abiotic, with the mixed community, with acetogens alone, with methanogens alone). To calculate electron recoveries we consider: 2 mM electron equivalents / eeq per mol H_2_ (according to: 2e^−^ + 2H^+^ → H_2_) and 8 mM eeq for each mol of methane or acetate (according to: CO_2_ + 8e^−^ + 8H^+^ → CH_4_ + 2H_2_O; and 2CO_2_ + 8e^−^ + 8H^+^ C_2_H_4_O_2_ + 2H_2_O). All experiments are run in triplicates (n=3).

**Figure 5.**
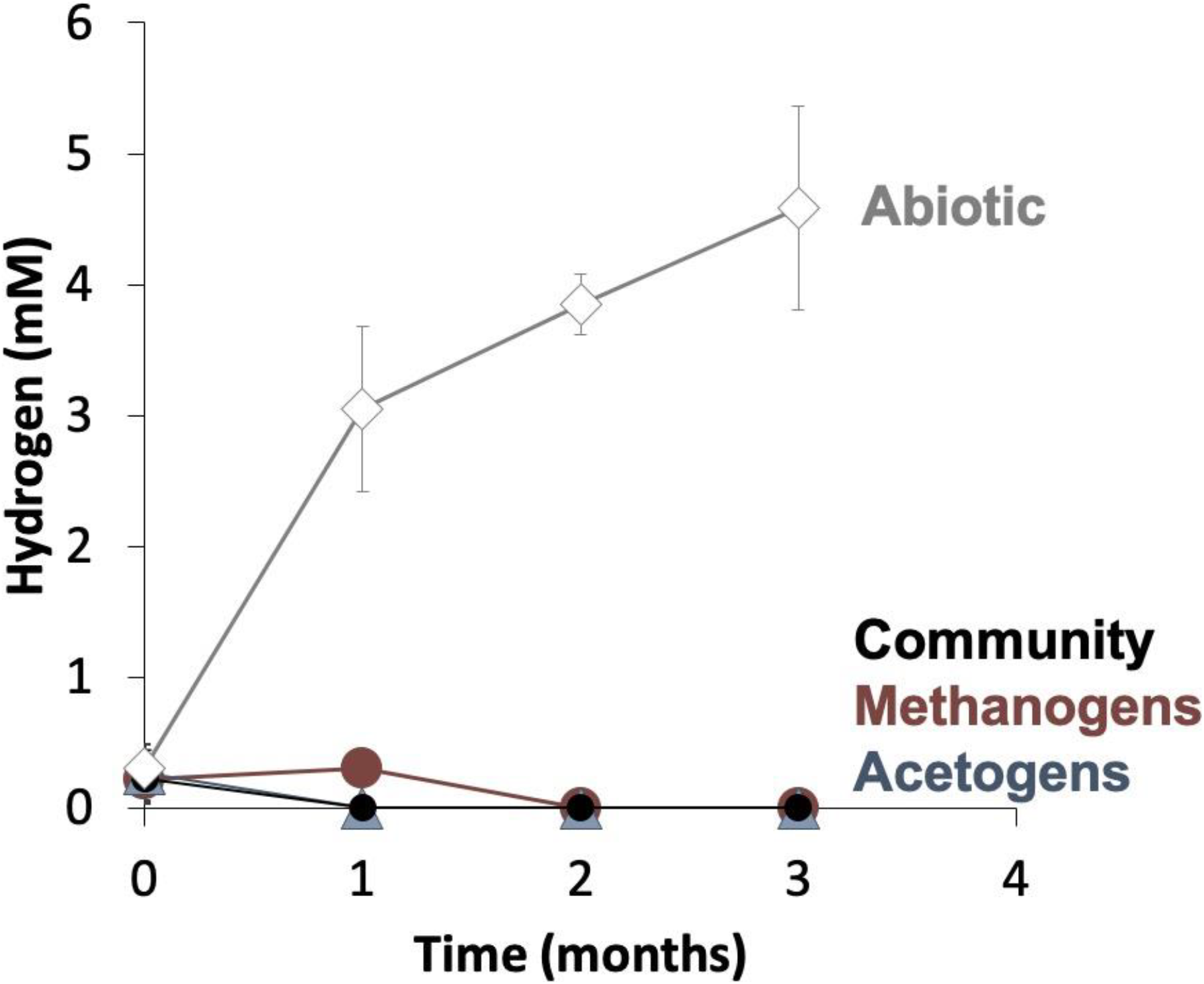
Hydrogen evolution in Fe^0^-media. Hydrogen production in abiotic controls (mineral media controls) with Fe^0^, versus incubations of cells (community, acetogens and methanogens alone) with Fe^0^. All experiments are run in triplicates (n=3) in parallel with experiments in Figure 3.

### 3.2 Unravelling interspecies interactions on Fe^0^

In all our enrichments, methane is the final product of the microbial community provided with Fe^0^ as electron donor and CO_2_ as electron acceptor. Thus, methanogens must interact with acetogens for example by consuming their metabolic product. There are three possible interspecies interactions that would give off only methane:

a. mutualism (+/+) when methanogens feed on the product of the acetogen both partners being influenced favorably one (acetogen) by the removal of metabolic product the other (methanogen) by the availability of a food substrate;
b. commensalism (0/+) when acetogens are unaffected by the presence of the methanogen, whereas methanogens are influenced favorably e.g., by feeding on acetate, the product of acetogenic metabolism.
c. parasitism/scavenging opportunism (−/+) when acetogens are negatively influenced by the presence of the methanogen, while methanogens are influenced favorably by the presence of the acetogen.

To test these scenarios, we carried out inhibition experiments to specifically block each metabolic group. Archaea methanogens were inhibited with 2-bromoethane sulfonate (BES), a coenzyme A analogue, resulting in favorable conditions solely for acetogens. Bacteria (including acetogens) were inhibited by a cocktail of antibiotics (kanamycin and ampicillin), thus favoring only methanogens. Then we compared corrosion by each group alone by documenting corrosion of via gravimetric measurements and metabolite production (Fig. 2, Fig. 3).

Gravimetric measurements showed that even when separated, methanogens and acetogens remained significantly more corrosive compared to cell free controls (Fig. 3c, 3d). However, methanogens were only slightly less corrosive than the mixed community (ca. 5% less, n=3, p=0.18). Whereas, acetogens were significantly more corrosive alone than they were in the mixed community (ca. 16% more; n=3, p=0.03). These results indicate that electron uptake from Fe^0^ by acetogens was negatively affected by the presence of the methanogens. The observation that acetogens are negatively impacted by methanogens was also supported by examining acetate metabolism. The rate of acetate build-up was lowered by the presence of the methanogens. Thus, the bacterial community alone accumulated acetate faster (Fig. 3c, 23% rate increase, n=3, p=0.0003) than when acetogens were members of the mixed community, substantiating the gravimetric results above, which revealed that acetogens were more corrosive alone (Fig. 2). Together, these results reflect that acetogens are negatively impacted (-) by the presence of the methanogens.

Methanogens on the other hand, produced 3-fold less methane alone than within the mixed community (Fig. 3d, n=3, p=0.0002). During the first month of incubation, rates of methanogenesis were 2-fold faster when acetogens were metabolically active (Fig. 3b) than they were when methanogens were incubated alone (Fig. 3d). In the mixed community, rates of methanogenesis increased significantly when acetogens reached stationary, now becoming 4-fold faster than when methanogens were incubated alone on Fe^0^. The rates of methanogenesis could not be linked to acetate-utilization, since acetate consumption (28±7 μM/day) did not match methane production (62.5±5μM/day). Thus, methanogens appear to be significantly favored (+) by the presence of the acetogens but only minutely by the acetate released by acetogens. It appears that methanogens may be favored by the presence of biogenic-chemicals and enzymes released by acetogens during the exponential-growth phase and especially during the stationary phase.

Altogether, our results show that during Fe^0^-corrosion, acetogens were negatively (-) impacted by the presence of methanogens whereas methanogens were positively (+) impacted, demonstrating a parasitic/opportunistic (−/+) type of interaction between these two physiologic groups.

A possible negative effect on the acetogens could be due to the alkalinization of the media by protons being dislocated from solution during CO_2_ conversion to methane. During CO_2_ conversion by methanogens the pH often becomes alkaline (Xu et al., 2014). Consequently, we verified the pH change over time. Typically, our cultivation media has a pH of 7.06±0.02. However, after six months of incubation, four cultures incubated solely with Fe^0^ exhibited an alkaline pH of 8.47±0.06. To verify whether alkalinization was dependent on cellular activity we monitored pH changes in abiotic Fe^0^-media, versus Fe^0^-media with cells (transfer 10) over the course of 30 days. We noticed a significantly higher alkalinization in media with cells compared to media without cells starting with day 15 (Fig. 6). Typically, CO_2_ reducing acetogenesis is negatively affected by alkaline conditions (Mohanakrishna et al., 2015), which explains the sudden decrease in acetogenic activity after day 15. On the other hand, alkaline conditions inhibit acetotrophic methanogenesis, while favoring CO_2_ reducing methanogenesis (Phelps and Zeikus, 1984).

**Figure 6.**
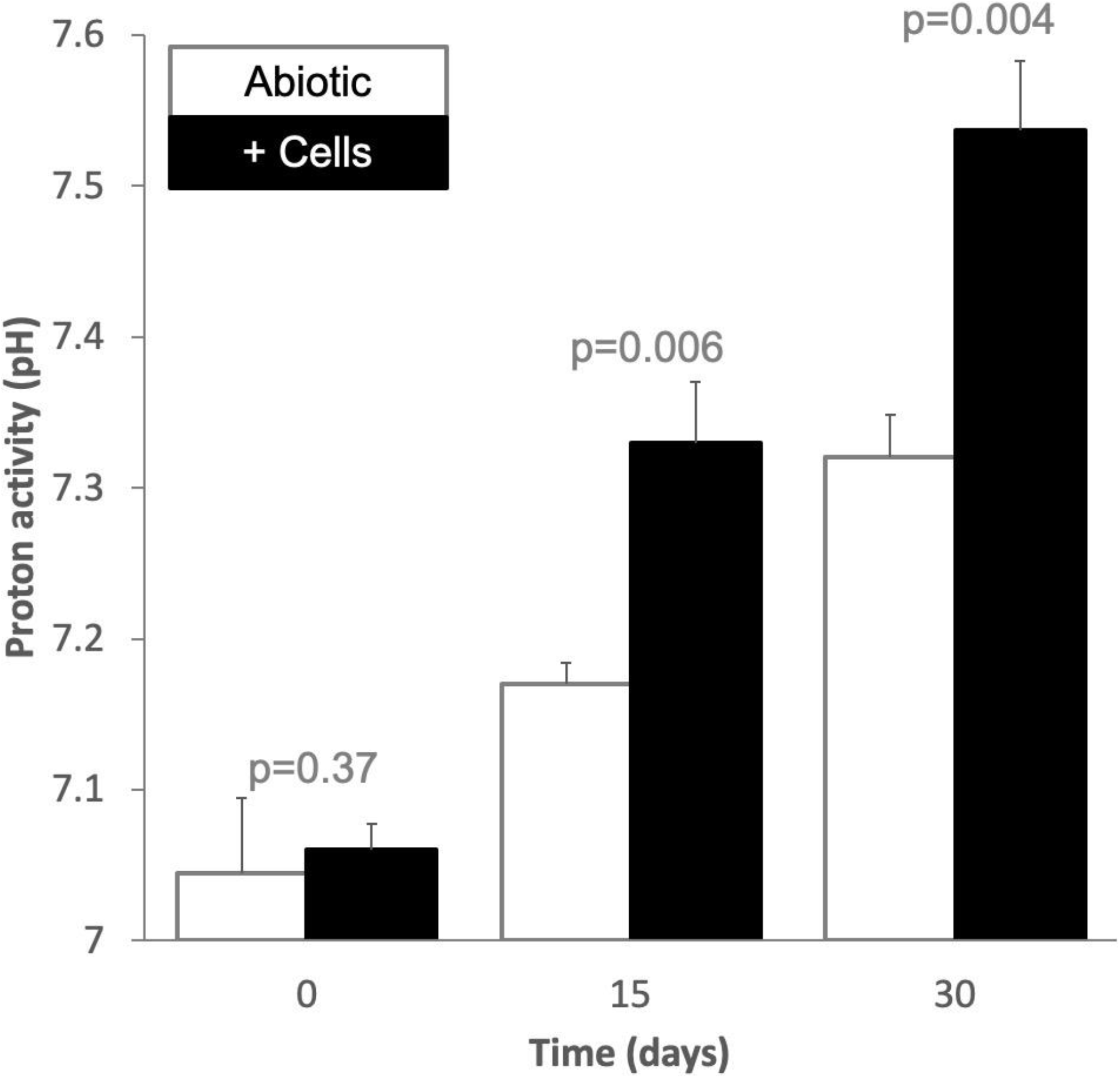
Proton activity (pH) changes during Fe^0^-incubations. pH changes in abiotic (white bars, mineral media controls) and cells (black bars, mixed corrosive community) when exposed to Fe^0^ for 1 month.

To further investigate the possible interplay between acetogens and methanogens we investigated the metabolic potential of the community by shotgun metagenomics.

### 3.3 Corrosive acetogens and methanogens

At transfer four, after 1 month of incubation, we studied the microbial community by shotgun metagenomics. Additional details about the results of shotgun metagenomics are available in the supplementary material. Pertinent to energy metabolism, the shotgun metagenome reinforced physiological observations confirming that the enriched corrosive community employs two major respiratory metabolisms: acetogenesis and methanogenesis.

#### 3.3.1 Acetogens

By shotgun metagenomics of the corrosive community, we discovered several genera of acetogens (Table 1). Of these genera, *Clostridium* had the best relative sequence abundance of all Bacteria (62.1%; Fig. 7a) and the best coverage of the Wood-Ljungdahl pathway for autotrophic acetogenesis (Fig.7, Table 1). *Clostridium* includes several species of autotrophs that use H_2_ as electron donor (Bengelsdorf et al., 2018) but also electrodes (Nevin et al., 2011) and Fe^0^ (Monroy et al., 2011). Besides, *Clostridium* was previously enriched on Fe^0^ from environments such as scraped bicycles thrown in a Dutch channel (Philips et al., 2019), rice paddies (Kato and Yumoto, 2015) or oilfield production waters (Ma et al., 2019). These studies along with ours suggests *Clostridium* is a likely corrosive acetogen when Fe^0^ becomes available in their environment.

**Table 1.** Gene coverage of the Wood Ljungdahl pathway in a shotgun metagenome from a climate lake. Here we only show genera of lithoautotrophic acetogenes.

| Genus | Gene coverage of the Wood Ljungdahl pathway* | Described lithoautotrophic acetogens of the genus |
| --- | --- | --- |
| *Clostridium* ** | 9 genes (*fhs*, *folD*, *metF*, *acsB*, *acsCD*, *cooS*, *pta*, *ack*) | 12 species (*C. aceticum*, *C. autoethanogenum*, *C. carboxidivorans*, *C. coskatii*, *C. difficile*, *C. drakei*, *C. formicoaceticum*, *C. ljungdahlii*, *C. magnum*, *C. methoxybenzovorans*, *C. ragsdalei*, *C. scatologenes*) |
| *Eubacterium* ** | 7 genes (*fhs*, *folD*, *acsB*, *acsD*, *cooS*, *pta*, *ack*) | 2 species (*E. aggregans*, *E. limosum*) |
| *Carboxydotherrmus* | 2 genes (*fhs*, *folD*) | 3 species (*C. ferrireducens*, *C. hydrogenoformans*, *C. pertinax*) |
| *Moorella* | 2 genes (*fhs*, *ack*) | 4 species (*M. glycerini*, *M. mulderi*, *M. thermoacetica*, *M. thermoautotrophica*) |
| *Treponema* | 2 genes (*fhs*, *ack*) | 1 species (*T. primitia*) |
| *Desulfotomaculum* ** | 2 genes (*metF*, *ack*) | 1 species (*D. thermobenzoicum*) |
| *Thermoanaerobacter* | 1 gene (*ack*) | 1 species (*T. kivui*) |

\*The Wood Ljungdahl pathway includes enzymes encoded by 13 genes which catalyze 10 reactions (see Fig. 7c): *fdhAB* (formate dehydrogenase), *fhs* (formate—tetrahydrofolate ligase), *folD* (methenyltetrahydrofolate cyclohydrolase), *metF* (methylenetetrahydrofolate reductase), *acsE* (5-methyltetrahydrofolate:corrinoid/iron-sulfur protein Co-methyltransferase), *acsAB* (CO-methylating acetyl-CoA synthase), *acsCD* (5-methyltetrahydrodarsinapterin:corrinoid/iron-sulfur protein CO-methyltransferase), *acsA/cooS* (Ni CO dehydrogenase), *pta* (phosphate acetyltransferase), *ack* (acetate kinase).
\*\* The 16S rRNA gene was detected in *Clostridium*, *Eubacterium* and *Desulfotomaculum*

**Figure 7.**
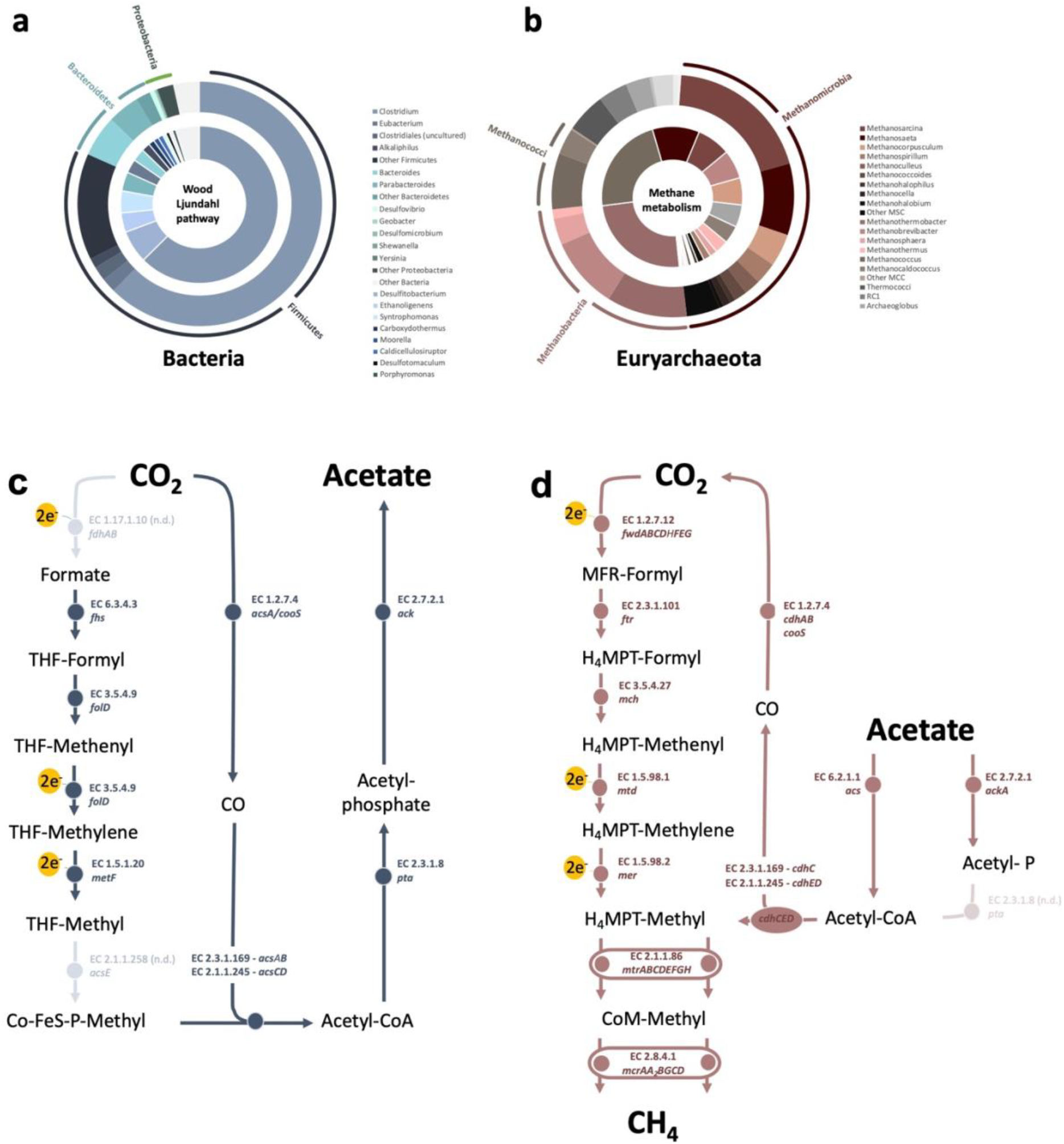
Microbes and pathways involved in Fe^0^ corrosion from a climate lake enrichment. Relative abundance of Bacteria (a) and Archaea (b) genera in a shotgun metagenome (outer circles) obtained from an SDU-pond community grown on Fe^0^ for 4 successive transfers. Inner circles are the relative distribution of genera according to genes of the Wood-Ljundgahl pathway for acetogenesis (a-inner circle) and genes for the methanogenesis pathway (b-inner circle). **Genes encoding for enzymes involved in C-modification during (c) acetogenesis and (d) methanogenesis.** The Wood Ljungdahl pathway involves: *fdh-*operon (formate dehydrogenase), *fhs* (formate—tetrahydrofolate ligase), *folD* (methenyltetrahydrofolate cyclohydrolase), *metF* (methylenetetrahydrofolate reductase), *acsE* (5-methyltetrahydrofolate:corrinoid/iron-sulfur protein Co-methyltransferase), *acs* (CO-methylating acetyl-CoA synthase), *acsCD* (5-methyltetrahydrosarcinapterin:corrinoid/iron-sulfur protein CO-methyltransferase), *acsA/cooS* (Ni CO dehydrogenase), pta (phosphate acetyltransferase), ack (acetate kinase). Methanogenesis involves: *fmd/fwd-*operon (formylmethanofuran dehydrogenase), *ftr* (formate—tetrahydrofolate ligase), *mch* (N^5^,N^10^-methenyltetrahydromethanopterin cyclohydrolase), *mtd* (F_420_-dependent N^5^, N^10^-methylene-H4MPT dehydrogenase), *mer* (N^5^,N^10^-methylene-H4MPT reductase), *mtr-*operon (coenzyme M methyltransferase), *mcr-*operon (methyl:coenzyme M reductase), *hdrA_1_B_1_C_1_* (ferredoxin:CoB—CoM heterosulfide reductase), *mvhAGD (-[NiFe]-hydrogenase*, *eha-*operon (energy-converting hydrogenase A), *ehb-*operon (energy-converting hydrogenase B), *frhABG* (F_420_-reducing hydrogenase) for CO_2_-reductive methanogenesis. Plus, *cdhED* (acetyl-CoA decarbonylase), *cdhC* (CO-methylating acetyl-CoA synthase), *acs* (acetyl-CoA synthetase), *ackA* (acetate kinase), *pta* (phosphotransacetylase) for accetotrophic methanogenesis.

Another autotrophic acetogen reasonably well represented in the shotgun metagenome was *Eubacterium* (1.9% of all Bacteria; Fig. 7a), which displayed the second-best gene recovery of the Wood Ljungdahl pathway (Table 1). *Eubacterium* includes two known species of autotrophic acetogens that use H_2_ as electron donor (Mechichi et al., 1998; Schwartz and Friedrich, 2006). Besides *Eubacterium* has been found in communities from thermogenic gas wells (Struchtemeyer et al., 2011) and stimulated Fe^0^ corrosion when co-existing with sulfate reducers, but have not been shown to be involved in Fe^0^ corrosion when alone (Dowling et al., 1992).

Other known autotrophic acetogens like *Moorella, Carboxydothermus*, *Treponema*, *Desulfotomaculum* and *Thermoanaerobacter* were scarce in this corrosive metagenome (<0.5% of all Bacteria) with a poor representation of the WL-pathway (Fig. 7a, Table 1). Of these, members of the genus *Desulfotomaculum* have been often shown to corrode Fe^0^ under sulfate reducing conditions (Anandkumar et al., 2009; Cetin and Aksu, 2009; Liu et al., 2019), but it remains to be investigated whether they can corrode Fe^0^ under acetogenic conditions.

Moreover, the metagenomes contained genera that include H_2_-utilizing species like *Bacillus* (1.6%), *Ruminococcus* (1.1%), *Desulfitobacterium* (1.1%) (Schwartz and Friedrich, 2006) and *Alkaliphilus* (1.3%) (Roh et al., 2007). These organisms use H_2_ when electron acceptors are available. But it is unknown whether any of these species can act as autotrophic acetogens, which is the only possibility in this enrichment. Of these, some *Bacillus* induced corrosion under nitrate reducing conditions (Wan et al., 2018; Xu et al., 2013), but in other instances *Bacillus* species inhibited corrosion by forming a passivating film of extracellular polymeric substances and/or calcite (Guo et al., 2019; Li et al., 2019). Corrosion was also inhibited by a *Desulfitobacterium* that oxidized Fe(III)-minerals to vivanite, forming a protective passivating layer on the metal surface (Comensoli et al., 2017). *Desulfitobacterium’s* protective effect against corrosion was even confirmed on archaeological iron artefacts (Comensoli et al., 2017).

Other well represented bacterial groups were *Bacteroidetes* (5.4% of all Bacteria) and *Parabacteroidetes* (4.2%), which are typically chemoorganotrophs (Sakamoto and Benno, 2006; Wexler, 2007) and may be involved in fermentation of decaying organic matter.

These results suggest that *Clostridium* is the central lithotrophic acetogen corroding Fe^0^ in these enrichments.

#### 3.3.2 Methanogens

The archaeal community was diverse with the highest relative sequence abundance matching members of the classes *Methanomicrobia* (45% of Archaea), *Methanobacteria* (24% of Archaea) and *Methanococci* (10% of Archaea).

The metagenome contained both acetate-utilizing methanogens and H_2_/CO_2_ utilizing methanogens. Acetate-utilizing methanogens belonging to the order *Methanosarcinales* were the best represented order making up for 32% of Archaea sequences, with *Methanosarcina* (19% of Archaea) showing the highest relative sequence abundance followed by *Methanosaeta* (9% of Archaea). Both genera showed good coverage of the methanogenesis pathway, however we could only find the 16S rRNA gene of *Methanosaeta*.

Generally, the role of acetotrophic *Methanosarcinales* in corrosion is assumed to be syntrophic and not directly corrosive (Mand et al., 2014; Zhang et al., 2003). However, previous studies reported that *Methanosarcina* species are corrosive (Daniels et al., 1987), and get enriched on Fe^0^ from the marine environment (Palacios et al., 2019a) or oilfield production waters (Ma et al., 2019). Often, *Methanosarcina* are members of corrosive biofilms on steel pipelines from oil production facilities (Lahme et al., 2021; Vigneron et al., 2016), gas industry pipelines (Zhu et al., 2003) and sewer system pipelines (Cayford et al., 2017). Unlike *Methanosarcina*, *Methanosaeta* have never been reported as independently corrosive, although they easily colonize Fe^0^-coupons in enrichments from oil production facilities (Mand et al., 2014) and have been often found as members of biofilms on corroded pipelines from such facilities (Conlette et al., 2016; Lahme et al., 2021).

Best represented H_2_/CO_2_-utilizing methanogens in relative gene abundance were members of *Methanothermobacter* (10% of Archaea) and *Methanobrevibacter* (9 % of Archaea) (Fig. 7b). These were also the genera with the highest number of gene hits for the CO_2_ reduction pathway (Table 2). *Methanothermobacter* species have been previously shown capable of Fe^0^-corrosion in pure culture (e.g., *M. thermoautotrophicum* (Daniels et al., 1987; Karr, 2013) or *M. thermoflexus* (Mayhew et al., 2016)). On the other hand, a *Methanothermobacter* strain isolated from a corroded oil pipeline could not corrode Fe^0^ alone, and required partnering with other microorganisms (Davidova et al., 2012). Often, *Methanothermobacter* are enriched or live within biofilms built on corroded oil pipelines (Liang et al., 2014; Okoro et al., 2017), corroded sea-submerged railway lines (Usher et al., 2014b), or oil facility infrastructure (Duncan et al., 2009; Lenchi et al., 2013; Suarez et al., 2019). Recently it was proposed that members of *Methanobacteriales* (order that includes *Methanothermobacter*) lead to the buildup of a carbonate-rich passivating layer with protective / anticorrosive properties (in ‘t Zandt et al., 2019). A possible protective role of *Methanothermobacter* in steel corrosion remains, however, to be demonstrated experimentally.

**Table 2.**
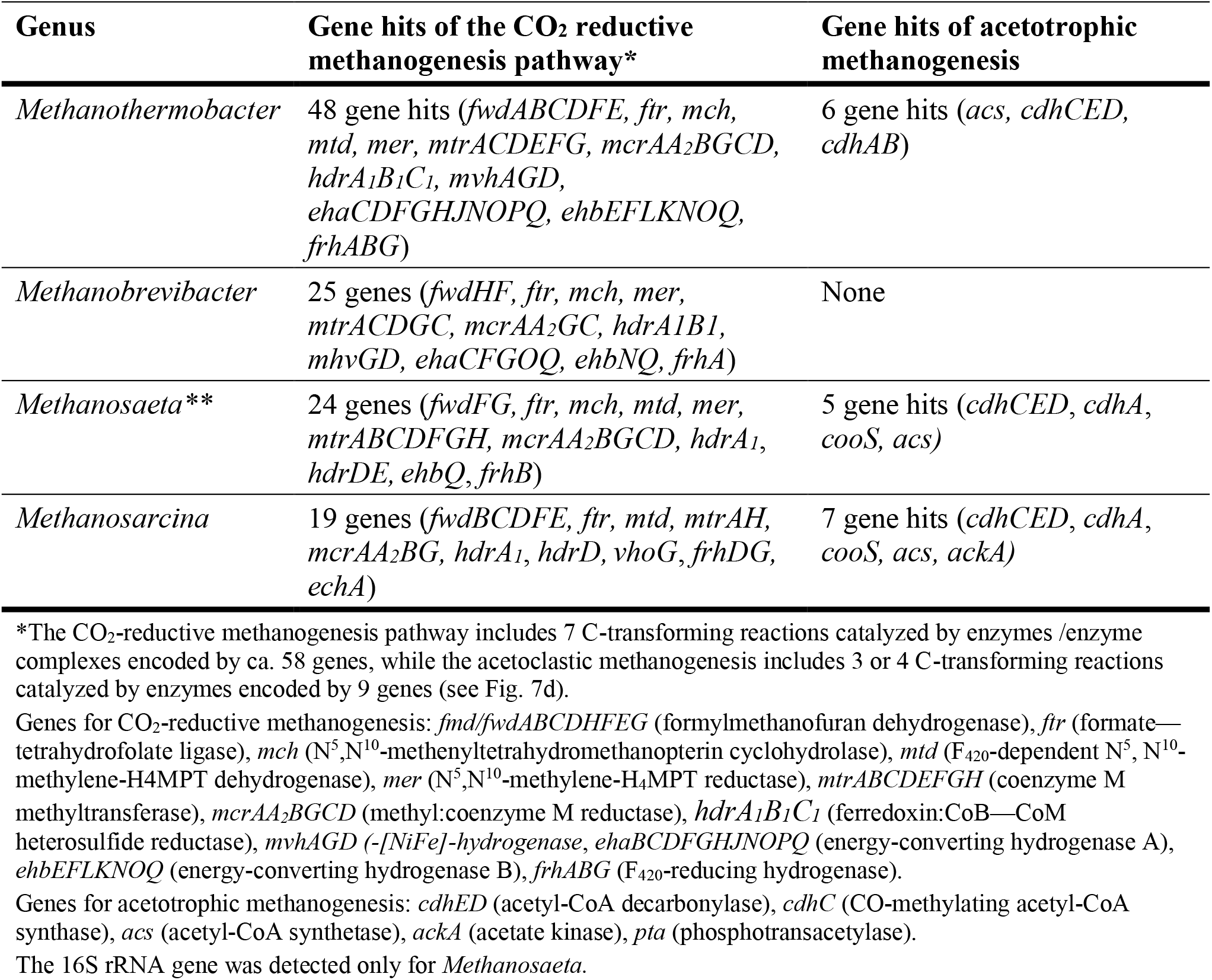
Gene hits of the CO_2_-reductive methanogenesis and acetotrophic methanogenesis pathway in a shotgun metagenome from a climate lake. Here we show genera that were relatively well represented.

*Methanobrevibacter* species on the other hand, are less often associated with corrosion with only a few studies reporting the presence of *Methanobrevibacter* in corrosive biofilms (Gomez-Alvarez et al., 2012; Schlegel et al., 2018; Usher et al., 2014b), while their role in corrosion remains unclear.

### 3.4 Possible mechanism of electron uptake during Fe^0^ corrosion

Since abiotic H_2_ does not explain the enhanced corrosion we observed, other mechanisms likely to enhance Fe^0^-corrosion by methanogens and acetogens in our enrichments such as: (1) the maintenance of low H_2_-partial pressure (Philips, 2020); (2) enzymatically mediated electron uptake (e.g., hydrogenases) (Deutzmann et al., 2015a; Lienemann et al., 2018b; Rouvre and Basseguy, 2016) or (3) direct electron uptake (e.g., via outermembrane cytochromes) (Tang et al., 2019).

In methanogens and acetogens it is unlikely that Fe^0^ oxidation proceeds via self-produced shuttles. This is because, although several other microorganisms produce their own shuttles to access extracellular electrons donors (Sund et al., 2007) acetogens and methanogens have never been documented to produce shuttles.

It is also unlikely that the community relies on a direct mechanism of electron uptake from Fe^0^, because it includes several groups of effective H_2_-utilizing microorganisms which could stimulate corrosion either by maintaining a low H_2_-partial pressure (Philips, 2020) or by enhancing electron uptake from Fe^0^ with the help of enzymes (Deutzmann et al., 2015b).

The first mechanism involves the maintenance of low H_2_ partial pressure at the Fe^0^ surface due to the low H_2_ thresholds of the microorganisms (concentration at which H_2_-uptake is not observed). It has been hypothesized that microorganisms with low H_2_-thresholds increase the H_2_ evolution reaction at the Fe^0^ surface (Philips, 2020). If microorganisms with low H_2_ thresholds were better Fe^0^ corroders in this corrosive community, strict hydrogenotrophic methanogens should be favored since it is the group with the lowest H_2_-threshold. They were not favored. Instead acetogens known to have high H_2_-thresholds were favored (Fig. 2, p=0.003). Generally, strict CO_2_-reducing methanogens exhibit very low H_2_-uptake thresholds (0.1-4 Pa, except for *M. wolfei* with 15 Pa) (Neubeck et al., 2016) followed by *Methanosarcinales* methanogens which exhibit higher H_2_-thresholds (13-46 Pa) during CO_2_ reductive methanogenesis. Even during organotrophy (acetate-grown), *Methanosarcinales* maintain a leveled H_2_-threshold concentration (20 Pa for *M. acetivorans* and 50 Pa for *M. thermophila*). On the other hand, CO_2_ reducing acetogens exhibit high H_2_-uptake thresholds (5-600 Pa, rarely below 10 Pa) (Philips, 2020). Thus, it is unlikely the maintenance of low H_2_ partial pressure plays aa major role in enhancing Fe^0^ corrosion in this community.

On the other hand, it is possible that H_2_-evolving enzymes boost microbial uptake of electrons from Fe^0^. Several H_2_-evolving enzymes were found in the metagenome of the corrosive community including: [FeFe] hydrogenases of acetogens, [NiFe] hydrogenases of strict hydrogenotrophic methanogens the formate/H_2_-evolving enzyme supercomplex of strict hydrogenotrophic methanogens (Mvh/Hdr: methyl viologen reducing hydrogenase / heterodisulfide reductase), but also enzymes that evolve H_2_ as a side reaction, like nitrogenases (Table 3).

**Table 3.**
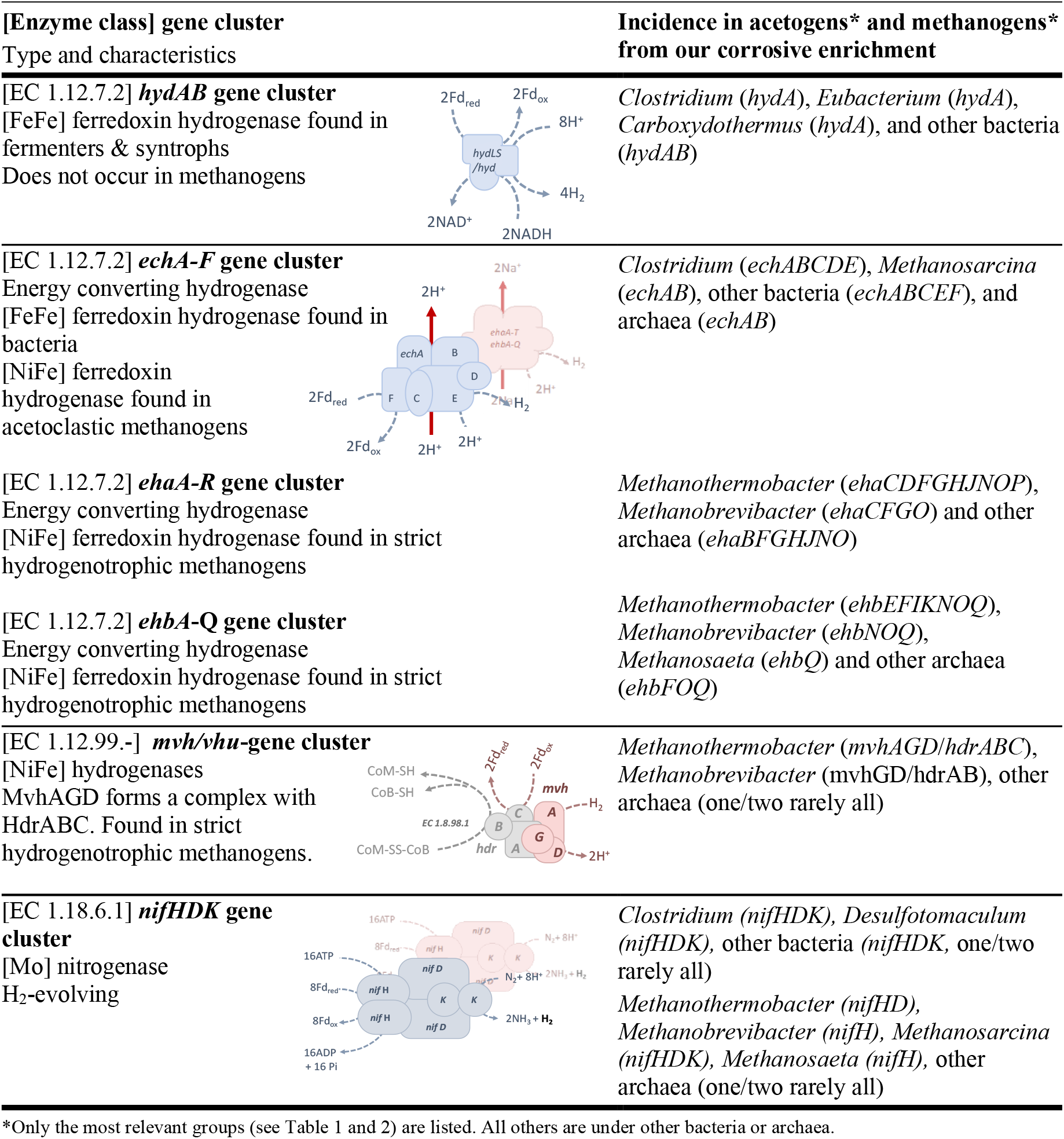
Gene hits for H_2_-evolving enzymes in a shotgun metagenome of a corrosive enrichment from a climate lake.

| [Enzyme class] gene cluster |  | Incidence in acetogens* and methanogens* |
| --- | --- | --- |
| Type and characteristics |  | from our corrosive enrichment |
| [EC 1.12.7.2] **hydAB gene cluster**<br>[FeFe] ferredoxin hydrogenase found in fermenters & syntrophs<br>Does not occur in methanogens |  | *Clostridium* (*hydA*), *Eubacterium* (*hydA*), *Carboxydotherrmus* (*hydA*), and other bacteria (*hydAB*) |
| [EC 1.12.7.2] **echA-F gene cluster**<br>Energy converting hydrogenase<br>[FeFe] ferredoxin hydrogenase found in bacteria<br>[NiFe] ferredoxin hydrogenase found in acetoclastic methanogens |  | *Clostridium* (*echABCDE*), *Methanosarcina* (*echAB*), other bacteria (*echABCEF*), and archaea (*echAB*) |
| [EC 1.12.7.2] **ehaA-R gene cluster**<br>Energy converting hydrogenase<br>[NiFe] ferredoxin hydrogenase found in strict hydrogenotrophic methanogens |  | *Methanothermobacter* (*ehaCDFGHJNOP*), *Methanobrevibacter* (*ehaCFGO*) and other archaea (*ehaBFGHJNO*) |
| [EC 1.12.7.2] **ehbA-Q gene cluster**<br>Energy converting hydrogenase<br>[NiFe] ferredoxin hydrogenase found in strict hydrogenotrophic methanogens |  | *Methanothermobacter* (*ehbEFIKNOQ*), *Methanobrevibacter* (*ehbNOQ*), *Methanosaeta* (*ehbQ*) and other archaea (*ehbFOQ*) |
| [EC 1.12.99.-] **mvh/vhu-gene cluster**<br>[NiFe] hydrogenases<br>MvhAGD forms a complex with HdrABC. Found in strict hydrogenotrophic methanogens. |  | *Methanothermobacter* (*mvhAGD/hdrABC*), *Methanobrevibacter* (*mvhGD/hdrAB*), other archaea (one/two rarely all) |
| [EC 1.18.6.1] **nifHDK gene cluster**<br>[Mo] nitrogenase<br>H<sub>2</sub>-evolving |  | *Clostridium* (*nifHDK*), *Desulfotomaculum* (*nifHDK*), other bacteria (*nifHDK*, one/two rarely all)<br>*Methanothermobacter* (*nifHD*), *Methanobrevibacter* (*nifH*), *Methanosarcina* (*nifHDK*), *Methanosaeta* (*nifH*), other archaea (one/two rarely all) |

\*Only the most relevant groups (see Table 1 and 2) are listed. All others are under other bacteria or archaea.

The [FeFe]-hydrogenases of acetogens have characteristic high H_2_-production rates (Adams, 1990) and can take up electrons directly from Fe^0^ to carry out proton reduction to H_2_ (Mehanna et al., 2008; Rouvre and Basseguy, 2016). The formate/H2-evolving enzyme supercomplex, Mvh/Hdr was shown to take up electrons directly from Fe^0^ to oxidize during formate synthesis (Lienemann et al., 2018a). In this case, formate-utilizing microorganisms would benefit. The [NiFe] hydrogenases of methanogens, especially those located on the MIC-island, appear to be crucial in inducing severe corrosion by hydrogenotrophic methanogens like *Methanobacterium* and *Methanococcus* (Lahme et al., 2021; Tsurumaru et al., 2018). However, the MIC island has not been identified in any other genera of methanogens.Thus, we could not detect the MIC-island in any of the genomes available for *Methanothermobacter, Methanobrevibacter* or *Methanosarcina* and *Methanosaeta* when using *in silico* PCR with specific MIC-island primers (Lahme et al., 2021).. On the other hand, using the same *in silico* approach we could for example detect the MIC-island in a *Methanobacterium* species (*M. congolense*) or in *Methanococcus* species (*M. maripaludis* OS7 and KA1) which have been documented to induce severe corrosion (Tsurumaru et al., 2018; Uchiyama et al., 2010).

Without the MIC-island these climate lake methanogens may be using an alternative approach to access Fe^0^-electrons effectively. Since acetate alone cannot explain the high methanogenesis rates, we hypothesize that acetogenic enzymes promote electron uptake from Fe^0^ to form H_2_ benefiting the methanogens. Thus, when acetogenesis becomes inhibited by the pH increase (prompted by methanogens delocalizing protons from solution into the gas phase), acetogen cell lysis cold release [FeFe]-hydrogenases and nitrogenases, and thereby produce H_2_ at the Fe^0^ surface. Enzymatically released H_2_ would benefit hydrogenotrophic methanogens like *Methanothermobacter* and *Methanobrevibacter* and could explain the acceleration of methanogenesis when acetogenesis ceased (Fig.3, Fig. 4). Regarding *Methanosarcinales,* their role in Fe^0^ corrosion could be simply that of commensals feeding on the acetate generated by acetogens, or they could be involved in direct electron uptake from extracellular electron donors (Rotaru et al., 2014b, 2014a; Yee et al., 2019; Yee and Rotaru, 2020) including Fe^0^ (Palacios et al., 2019a). However, direct electron uptake in *Methanosarcinales* is poorly understood and remains to be further investigated.

## 4 Conclusion

Interspecies interactions at the Fe^0^-surface have been poorly investigated. Here we bring evidence for an opportunistic interaction (+/−) on Fe^0^ between methanogens (*Methanothermobacter*, *Methanobrevibacter* and *Methanosarcinales*) and acetogens (esp. *Clostridium*) enriched from the sediments of a climate lake. We observed that the acetogens were more effective Fe^0^-corroders (and acetate producers) when decoupled from methanogens with the help of a specific inhibitor – 2-bromoethane sulfonate. Methanogens, on the other hand, became less effective Fe^0^-dependent methane producers when decoupled from acetogens with the help of antibiotics. Since abiotic H_2_ could not explain the corrosion rates, we screened a shotgun metagenome for hints of enzymatic-mediated H_2_-evolution. The metagenome of acetogens (esp. *Clostridium*) contained [FeFe] hydrogenases, effective in direct electron uptake from Fe^0^ in order to reduce protons to H_2_. On the other hand, the MIC-island, a typical indicator for enzymatic mediated electron uptake in methanogens was not detected in *Methanothermobacter*, *Methanobrevibacter* or *Methanosarcinales*. As acetogenesis ceased methanogenesis rates picked up, the later could not be explained by acetate turnover. Therefore, we propose that the methanogenic community benefits from H_2_-evolving enzymes released by acetogens during stationary phase. With these results we expand the diversity of microbial interactions on Fe^0^ between acetogens and methanogens (from competition (Palacios et al., 2019b) to opportunism (this study)). Some studied suggested that methanogens like *Methanothermobacter* may have a protective role during corrosion (in ‘t Zandt et al., 2019), however here we show how such methanogens are positively affected by acetogenic partners. In consequence we recommend that the corrosive effects of methanogens should be investigated not only in pure culture but also in consortia with bacterial partners before suggesting they have a protective anticorrosive role.

## 6 Author Contributions

AER and PAP developed the idea. PAP carried out all experimental work and some of the analyses. AER carried out some of the data analyses. PAP drafted the paper. AER wrote the final manuscript.

## 8 Acknowledgments

We would like to thank Oona Snoeyenbos-West, Carolin Löscher, Satoshi Kawaichi and Christian Furbo Reeder for help with sampling and valuable discussions. We would like to thank Joy Ward for help with preparing samples for scanning electron microscopy and we’d like to recognize the University of Massachusetts electron microscopy facility which provided access and training of PAP on the FEI Magellan XHR-SEM.

